# FusOn-pLM: A Fusion Oncoprotein-Specific Language Model via Focused Probabilistic Masking

**DOI:** 10.1101/2024.06.03.597245

**Authors:** Sophia Vincoff, Shrey Goel, Kseniia Kholina, Rishab Pulugurta, Pranay Vure, Pranam Chatterjee

## Abstract

Fusion oncoproteins, a class of chimeric proteins arising from chromosomal translocations, drive and sustain various cancers, particularly those impacting children. Unfortunately, due to their intrinsically disordered nature, large size, and lack of well-defined, druggable pockets, they have been historically challenging to target therapeutically: neither small molecule-based methods nor structure-based approaches for binder design are strong options for this class of molecules. Recently, protein language models (pLMs) have demonstrated success at representing protein sequences with information-rich embeddings, enabling downstream design applications from sequence alone. However, no current pLM has been trained on fusion oncoprotein sequences and thus may not produce optimal representations for these proteins. In this work, we introduce **FusOn-pLM**, a novel pLM that fine-tunes the state-of-the-art ESM-2 model on fusion oncoprotein sequences. We specifically introduce a novel masked language modeling (MLM) strategy, employing a binding-site probability predictor to focus masking on key amino acid residues, thereby generating more optimal fusion oncoprotein-aware embeddings. Our model improves performance on both fusion oncoprotein-specific benchmarks and disorder prediction tasks in comparison to baseline ESM-2 representations, as well as manually-constructed biophysical embeddings, motivating downstream usage of FusOn-pLM embeddings for therapeutic design tasks targeting these fusions. We have made our model publicly available to the community at https://huggingface.co/ChatterjeeLab/FusOn-pLM.

## Introduction

Fusion oncoproteins arise from chromosomal rearrangements that fuse segments of two distinct genes. The resulting mutants contain unrelated functional domains connected by long regions of disorder. This flexible configuration promotes constitutive activation or aberrant regulation of the fusion proteins, driving oncogenic transformation and tumor development [Angione et al., 2021]. Thousands of unique fusion oncoproteins have been discovered by sequencing patient tumors, and several common culprits such as EWS::FLI1 in Ewing’s sarcoma [Delattre et al., 1992], PAX3::FOXO1 in alveolar rhabdomyosarcoma (ARMS) [Linardic, 2008], SS18::SSX1 in synovial sarcoma [McBride et al., 2018], and EML4::ALK proteins in non-small-cell lung cancer [Soda et al., 2007] are well characterized in the literature. However, even the best understood fusion oncoproteins have proven to be elusive drug targets due to their structural instability and absence of defined binding pockets [Tripathi et al., 2023] (Figure 1A). For small molecules that are able to bind fusion oncoproteins, for example EWS::FLI1, these compounds do not achieve strict fusion specificity, binding to one of their head or tail protein counterparts that are often critical transcription factors for cellular homeostasis [Erkizan et al., 2009, Vital et al., 2023]. As such, biologics, such as antibodies, miniproteins, and peptides, represent attractive therapeutic alternatives, but necessitate advanced design approaches for specific targeting to these undruggable proteins.

**Figure 1.**
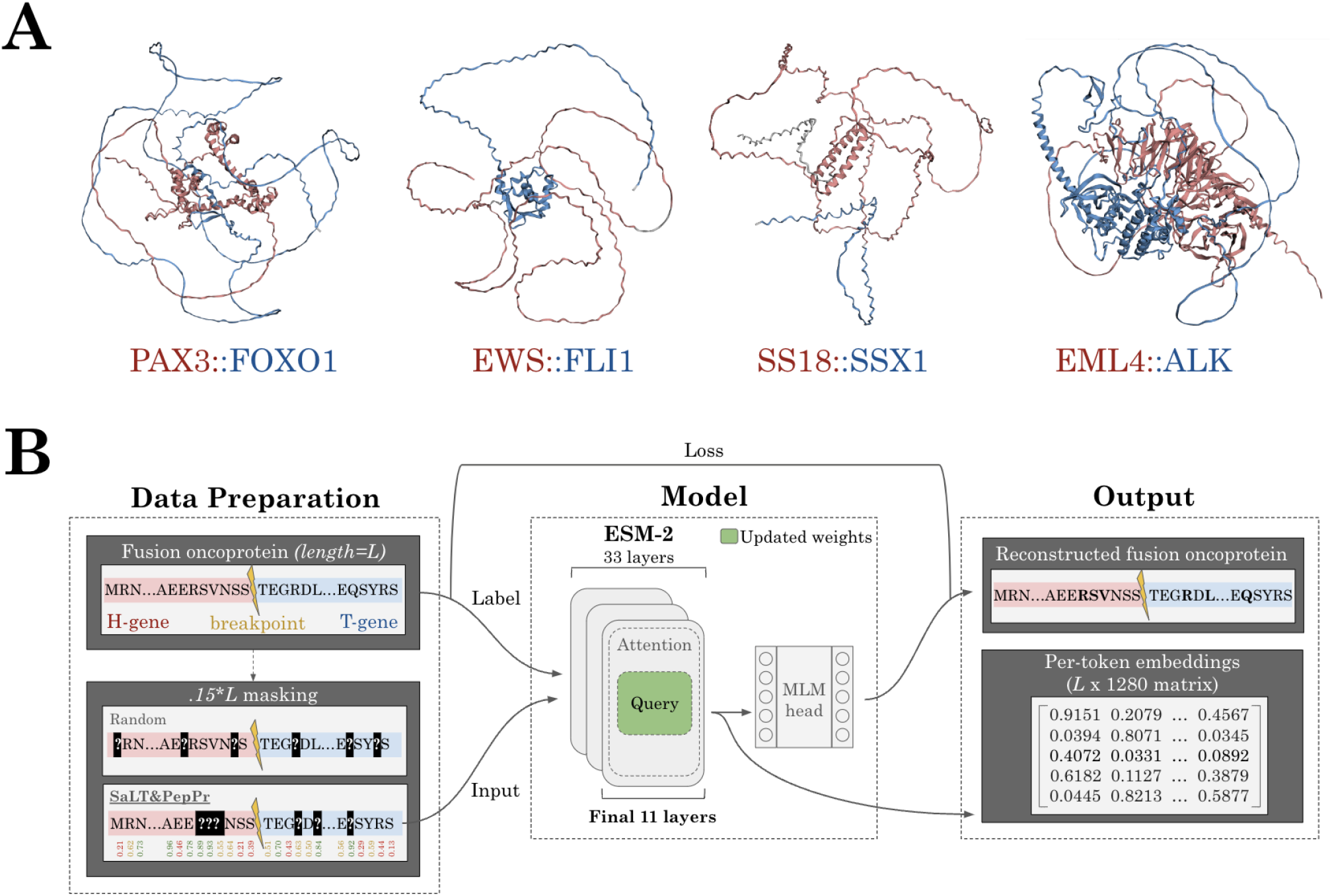
Overview of FusOn-pLM. **A** AlphaFold3 structures of four well-studied fusion oncoproteins. Fusion protein segments derived from the head gene are colored in red; tail in blue. **B** FusOn-pLM pipeline. **Data preparation**: Fusion oncoprotein sequences (length *L*) undergo 15% masking by either: (1) random masking, where each amino acid has equal likelihood of selection, or (2) SaLT&PepPr-based masking, where potential binding sites on the fusion oncoprotein are more likely to be masked. SaLT&PepPr-based masking produced the final FusOn-pLM model. The masked sequence is fed as input and the original sequence as label into the **model**: 33-layer ESM-2-650M with an MLM head. In the top third of the model (final eleven layers), the query weights are unfrozen for fine-tuning. **Output**: the MLM head outputs an attempted reconstruction of the original sequence, which is compared with the label to calculate loss. FusOn-pLM embeddings, of shape *[L, 1280]*, are extracted from the final layer of the ESM-2-650M encoder stack.

Recently, structure-based prediction and design models, such as AlphaFold and RFDiffusion [Jumper et al., 2021, Abramson et al., 2024, Watson et al., 2023], have accelerated the design of biologics targeting pathogenic proteins. These tools, by default, fail to accurately capture the structure of numerous conformationally unstable proteins, limiting their usefulness for fusion oncoprotein targeting [Piovesan et al., 2022]. Meanwhile, protein language models (pLMs), such as ESM-2 and ProtT5, have been trained on the amino acid sequences of over 250 million proteins, from the exceedingly stable to the intrinsically disordered [Lin et al., 2023, Elnaggar et al., 2022]. They capture physicochemical, structural, and functional properties of proteins from their sequence alone, and have even been extended to design novel proteins [Ferruz et al., 2022, Madani et al., 2023] and binders [Brixi et al., 2023, Bhat et al., 2023, Chen et al., 2023]. However, these models were not trained on fusion oncoprotein sequences, which are functionally and structurally distinct from their wild-type counterparts due to their altered binding sites and unique breakpoint junctions.

To fill this critical gap, we fine-tune the state-of-the-art ESM-2 pLM on over 35,000 fusion oncoprotein sequences collected from the FusionPDB and FOdb databases [Kumar et al., 2024, Tripathi et al., 2023]. We specifically unfreeze the query weights and biases of the final eleven layers of the ESM-2 model and fine-tune these parameters via a masked language modeling (MLM) head (Figure 1B). To encourage our model to learn the distinct features of fusion oncoproteins responsible for their function, we introduce a novel probabilistic masking strategy. We apply our recent SaLT&PepPr model to predict and bias masking toward residues that are most likely to participate in protein-protein interactions (PPIs) [Brixi et al., 2023] (Figure 1B), and therefore likely to facilitate novel oncogenic interactions [Mukherjee et al., 2022]. Our results demonstrate that the output embeddings from our SaLT&PepPr-based masking strategy outperform baseline embeddings on diverse fusion oncoprotein-specific tasks as well as on standard disorder prediction benchmarks, while distinctly representing the fusion oncoproteins from their original head and tail protein counterparts. In total, these results motivate the application of our fusion-specific embeddings for therapeutic design tasks.

## Methods

### Model Training Set Curation

Model training data was curated from the FusionPDB and FOdb databases [Kumar et al., 2024, Tripathi et al., 2023]. Specifically, 41,420 FusionPDB and 4,536 FOdb unique amino acid sequences containing only the 20 natural amino acids were collected for downstream model training. Proteins longer than 2000 amino acids were removed due to GPU memory limits. 1,308 duplicates from database overlap were removed, and 177 FOdb sequences were held out for benchmarking tasks. All remaining sequences were clustered using MMSeqs2 easy clustering module with a minimum sequence identity threshold of 30% and a coverage threshold of 80% [Steinegger and Söding, 2017]. The resulting clusters were split at a 80/10/10 ratio into a training set (31,788 proteins, 79.8%), validation set (4,030 proteins, 10.1%), and testing set (4,013 proteins, 10.1%).

### Benchmarking Dataset Curation

Datasets for the three puncta-related benchmarking tasks were collected from FOdb [Tripathi et al., 2023]. 177 FOdb sequences were held out for three classification tasks concerning the tendency of fusion oncoproteins to form condensates (puncta) and the cellular localizations of these puncta. These sequences were clustered using MMSeqs2 easy clustering module with a minimum sequence identity threshold of 30% and a coverage threshold of 30% (larger coverage thresholds led to formation of very few clusters). For each task, the clusters were split at 80/20 ratio into train and test sets with similar ratios of class 0 to class 1. For puncta propensity of formation, there were 143 train sequences (80.8% of total; 35.7%-64.3% class 0-1) and 34 test sequences (19.2% of total; 35.3%-64.7% class 0-1). For puncta localization to the nucleus, there were 143 train sequences (80.8%; 59.4%-40.6% class 0-1) and 34 test sequences (19.2%; 58.8%-41.2% class 0-1). For puncta localization to the cytoplasm, there were 141 train sequences (79.7%; 64.5%-35.5% class 0-1) and 36 test sequences (20.3%; 63.9%-36.1% class 0-1).

The next benchmarking task involved predicting fusion oncoprotein disease outcomes. Cancer associations for the test set (4,013 proteins) were extracted from FusionPDB [Kumar et al., 2024]. This data was originally collected from The Cancer Genome Atlas (TCGA), which provided full definitions of each cancer acronym [Weinstein et al., 2013]. The top two cancer types were breast invasive carcinoma (BRCA, 583 sequences) and stomach adenocarcinoma (STAD, 489 sequences). Fusion oncoproteins causing these diseases were extracted and clustered using MMSeqs2 easy clustering module with a minimum sequence identity threshold of 30% and a coverage threshold of 80%. These clusters were split into train and test sets: 859 train (80.13%; 54.4%-45.6% BRCA-STAD), 213 test (19.87%; 54.5%-45.5% BRCA-STAD).

We utilized the dataset from [Lotthammer et al., 2024] for an additional test where models predicted ensemble dimensions of intrinsically disordered regions (IDRs). The dataset contains IDR sequences from both synthetic and naturally occurring proteins, as well as simulation-derived IDR properties including asphericity and radius of gyration. Radius of gyration represents the average distance between IDR amino acids and the protein’s center of mass. Asphericity is a dimensionless parameter ranging from 0 (sphere) to 1 (prolate ellipsoid) which describes the shape of an IDR ensemble. To prepare training data, the 47,114 IDR sequences were first clustered using the MMSeqs2 easy clustering module with a minimum sequence identity of 30%, a coverage threshold of 50%, coverage mode 1 (target coverage - recommended mode for fragments, which are abundant in the IDR dataset), and cluster mode 2 (greedy incremental - recommended pairing for target coverage). The resulting clusters were split at an 80/10/10 ratio into a training set (37,737 IDRs, 80.10%), validation set (4,665 proteins, 9.90%), and testing set (4,712 proteins, 10.00%).

For disorder prediction, 1,316 sequences were collected from DisProt [Aspromonte et al., 2023] and the RCSB PDB, all of which are evaluated in The **C**ritical **A**ssessment of Protein **I**ntrinsic **D**isorder Prediction (CAID) experiment [Necci et al., 2021]. For each sequence, per-residue annotations indicating disorderliness (class 1) or structure (class 0) were provided by CAID. These sequences and their associated per-residue labels of disorder were split using the MMSeqs2 easy clustering module with a minimum sequence identity of 30% and a coverage threshold of 80%. The resulting clusters were split at an 80/10/10 ratio into a training set (1,051 proteins, 79.8%), validation set (131 proteins, 10.0%), and testing set (134 proteins, 10.2%).

To extend our per-residue disorder analysis beyond the CAID dataset to our target class of interest, fusion oncoproteins, we assembled pseudo-labels for fusion oncoproteins based on pLDDT scores from AlphaFold2. We queried our test set (4,013) in FusionPDB Level III (75 hits) and collected all available AlphaFold2 .cif files (66). According to AlphaFold2, residues with pLDDTs < 50 were considered disordered, and residues with pLDDTs > 50 were considered structured. Following this guidance, we converted pLDDTs for each amino acid in each sequence into zeros (< 50) and ones (≥50) [Abramson et al., 2024].

### Amino Acid Masking Strategies

To force comprehension of physicochemical features of fusion oncoproteins, we employ a focused probabilistic masking strategy on input amino acid sequences. Specifically, we mask 15% of the full sequence, as this percentage has performed well in prior studies [Devlin et al., 2018]. Since fusion oncoproteins represent the interaction of two distinct proteins, we masked amino acids that are likely to participate in PPIs as determined by the output probabilities of SaLT&PepPr [Brixi et al., 2023], which predicts a per-amino acid probability of binding. Our masking strategy is as follows:

Let ***x*** = (*x*_1_, *x*_2_, … , *x*_*n*_) be the input amino acid sequence of length *n*, and *p*_*i*_ be the probability that the amino acid *x*_*i*_ participates in a PPI as predicted by SaLT&PepPr. Define *M* as the set of masked positions such that |*M* | = ⌈0.15*n*⌉.

We select *M* using the following probabilistic strategy:

1. Compute the binding probability for each amino acid from SaLT&PepPr: ***P*** = (*p*_1_, *p*_2_, … , *p*_*n*_).
2. Normalize the probabilities using the `softmax` function:

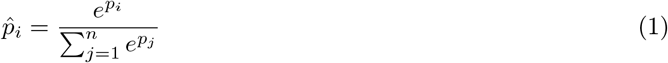

1. Sample *M* by selecting ⌈0.15*n*⌉ positions according to the normalized probabilities 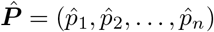 .Mathematically, the selection of *M* can be described as:

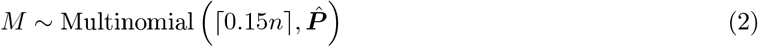

Alternatively, for the random 15% masking, we uniformly sample *M* from the set {1, 2, … , *n*} without replacement:

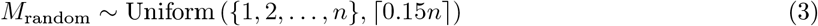

A visualization of the masking strategy is shown in Figure 1.

### Model Architecture and Training

FusOn-pLM is a fine-tuned encoder on curated fusion oncoprotein sequences trained via an MLM task to create fusion oncoprotein-aware embeddings (Figure 1). To preserve comprehension of wild-type proteins, we train FusOn-pLM with an MLM head on ESM-2-650M [Lin et al., 2023], where amino acid tokens (masked using the respective masking strategy) are passed into ESM-2-650M to retrieve their output embeddings. The MLM loss function ℒ_MLM_ is defined as:

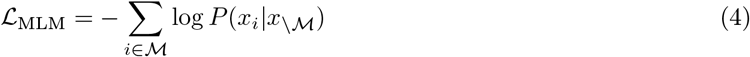

where ℳ represents the set of masked positions in the input sequence, *x*_*i*_ is the true amino acid token at position *i*, and *x*_\*ℳ*_ denotes the sequence with the masked tokens excluded.

FusOn-pLM was trained on one NVIDIA H100 GPU with 80 GB of VRAM each for 14 epochs with batch size of 8 and learning rate of 5e-5. The `Adam` optimizer was utilized with no weight decay. Only fusion oncoproteins of length 2000 or shorter were used for training; short sequences were padded to this maximal length.

To optimize performance while avoiding overfitting on our new sequences, we unfroze only the query weights in a fraction of ESM-2-650M layers and benchmarked the ensuing models at each epoch (Figure 1B). Using random masking, we trained models with a minimum of three and maximum of seventeen unfrozen terminal layers, to avoid sacrificing on batch size (Figure 1B).

### Fusion Oncoprotein Property Benchmarks

In recent works, certain fusion oncoproteins have been shown to form puncta via phase separation, a hallmark pathology preceding cancer phenotypes and tumor proliferation [Jiang et al., 2020]. To determine if our FusOn-pLM embeddings produce accurate numerical representations of fusion oncoproteins, we evaluated the embeddings’ performance on predicting the propensity of puncta formation as well as predicting if puncta form in the nucleus or cytoplasm. Here, we utilized 177 sequences from FOdb with experimental data on puncta formation for pLM embedding evaluation [Tripathi et al., 2023]. Cancer associations from FusionPDB were further used to evaluate FusOn-pLM’s ability to distinguish fusion proteins that drive different malignancies.

Puncta formation and localization predictions were treated as a binary class, where label 0 or 1 represented a lack or presence of puncta formation in a given area. For the cancer association task, two binary classes were defined for 1,072 test-set proteins: BRCA (class 0) and STAD (class 1). We compared FusOn-pLM embeddings against three others: 1) Base wild-type ESM-2-650M embeddings, 2) FOdb embeddings, which are 25 physicochemical features manually curated by FOdb for only these 177 proteins, and finally, 3) Basic one-hot embeddings. We leveraged the standard binary cross-entropy loss function and minimized this loss function for each task using the XGBoost model with 50 trees via `scikit-learn` [Buitinck et al., 2013]. The binary cross-entropy loss is defined as:

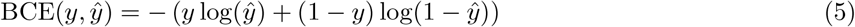

### Disorder Property Benchmark

Structural disorder is a hallmark feature of fusion oncoproteins. To assess FusOn-pLM’s resulting knowledge of disorder, we utilized FusOn-pLM to predict physical properties of IDRs, as defined previously [Lotthammer et al., 2024]. For each sequence, we computed three sets of embeddings: FusOn-pLM, ESM-2-650M, and one-hot. To evaluate each method, we utilized a linear probing technique in which embeddings were passed through a defined multi-layer perceptron (MLP) network with three fully connected layers. The input layer performed dimensionality reduction to hidden dimension 640 and passed the output through a ReLU activation function, followed by layer normalization and dropout regularization with a probability of 0.2. This structure was repeated for two more iterations, shrinking the hidden dimension to 320 and finally culminating in a single neuron: a scalar output to enable property prediction. Models were trained to minimize the mean square error (MSE). The MSE loss function is defined by:

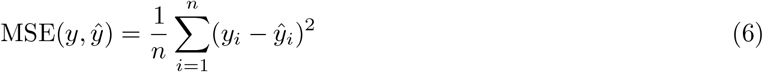

To identify optimal training parameters, we screened two batch sizes (32, 64) and various learning rates between 1e-5 and 1e-3. One model for each embedding was developed for each prediction variable (asphericity, *R*_*g*_), resulting in a total of 6 models. Ultimately, all models were trained with identical hyperparameters: 20 epochs of training with a batch size of 64 and learning rate 3e-4. Early stopping was implemented to prevent model overfitting. Each model was evaluated on a held-out test set by predicting each property given the sequence embedding alone. *R*^2^ scores were calculated to assess model performance. The true values and predicted values were plotted to visualize the model’s accuracy, with an ideal fit line included for reference.

### CAID Benchmark

To further assess FusOn-pLM’s understanding of disorder, we used disorder prediction at a per-residue basis as a benchmarking task. This test is formally defined by the CAID experiment, where a score is assigned to each residue for its propensity of being intrinsically disordered, thereby predicting which residues likely contribute to the protein’s disordered nature [Necci et al., 2021]. Each sequence used in the per-residue disorder prediction task was assigned a set of ground truth labels indicating if each residue was disordered (class 1) or structured (class 0). With defined binary classes, we minimized the standard binary cross-entropy loss, as defined in Equation (5), by training a single-layer perceptron classification head in PyTorch that outputs the probability that each residue in the sequence is either disordered (probability > 0.5, class 1) or structured (probability < 0.5, class 0). In this task, we compared the performance of FusOn-pLM embeddings against wild-type ESM-2-650M embeddings by utilizing each as input into the classification head. With predicted per-residue disorder labels for each embedding type, we visualized regions of predicted disorder on 3D structures of wildtype proteins (from the CAID dataset) and fusion oncoproteins (from our dataset) to assess the model’s efficacy in discerning between regions of structure and disorder and capturing this information in its embeddings.

### Embedding Exploration

To explore how FusOn-pLM embeddings capture the physicochemical properties of fusion oncoproteins, we first conducted a dimensionality reduction analysis on both fusion oncoprotein embeddings and/or their head and tail proteins using t-SNE via the `sklearn.manifold` module. We utilized low mutual information physicochemical features curated in FOdb as the basis for clustering based on embedding properties. The 2D FusOn-pLM t-SNE representations were finally plotted on scatter plots in `matplotlib`. For sequence property visualization, we adopted a color-coding scheme wherein each point in the scatter plot represents a fusion oncoprotein sequence and is colored according to its value for a specific physicochemical feature.

## Results

### Probabilistic masking enables focused training

First, we sought to identify which masking strategy obtains optimal fusion oncoprotein embeddings. Our training results demonstrate that while both SaLT&PepPr-based and random masking produced similar training results with low perplexity values (Table 1), optimal results on preliminary benchmarking were reached before the model converged or displayed evidence of overfitting, indicating that training loss alone cannot be relied upon to choose the final model. As such, our final, optimal model was trained with 11 unfrozen layers using SaLT&PepPr-based masking. By freezing the weights in the remaining 22 layers of ESM-2 and the random MLM head, we enable efficient adaptation to fusion oncoproteins with a small set of trainable parameters. In total, our final FusOn-pLM model consists of 651,163,541 parameters in the ESM-2 encoder stack (18,036,480 of which are trainable parameters) and 1,684,513 parameters in its MLM head.

**Table 1:** FusOn-pLM perplexities at different training stages with specified masking strategies.

| Masking | Epoch | Train pPL | Val pPL | Test pPL |
| --- | --- | --- | --- | --- |
| SaLT&PepPr 15% | 14 | 4.731 | 4.827 | 4.851 |
| SaLT&PepPr 15% | 20 | 4.455 | 4.598 | 4.607 |
| Random 15% | 14 | 4.620 | 4.700 | 4.840 |
| Random 15% | 20 | 4.342 | 4.475 | 4.506 |

### FusOn-pLM generates fusion oncoprotein-relevant representations

To determine if FusOn-pLM produces relevant embeddings, we sought to evaluate its performance on downstream fusion oncoprotein-specific tasks. We first assessed the embeddings’ ability to accurately predict the propensity and localization of puncta, critical formations driving cancer pathology [Tripathi et al., 2023]. From our classification metrics on puncta formation propensity, we demonstrate that FusOn-pLM embeddings strongly outperform ESM-2-650M, FOdb, and one-hot embeddings on all relevant classification metrics across the entire held-out test dataset (Figure 2A), which is also the case for predicting localization to the nucleus, the primary location of fusion oncoproteins [Angione et al., 2021] (Figure 2B). While FOdb embeddings perform strongly on cytoplasm localization prediction, FusOn-pLM proves most effective on the critical AUROC metric (Figure 2C), and comparatively outperforms all other embeddings for the prediction of carcinoma class (Figure 2D). In total, these results indicate that FusOn-pLM learns representations capturing key semantics and properties encoded in fusion oncoprotein sequences.

**Figure 2.**
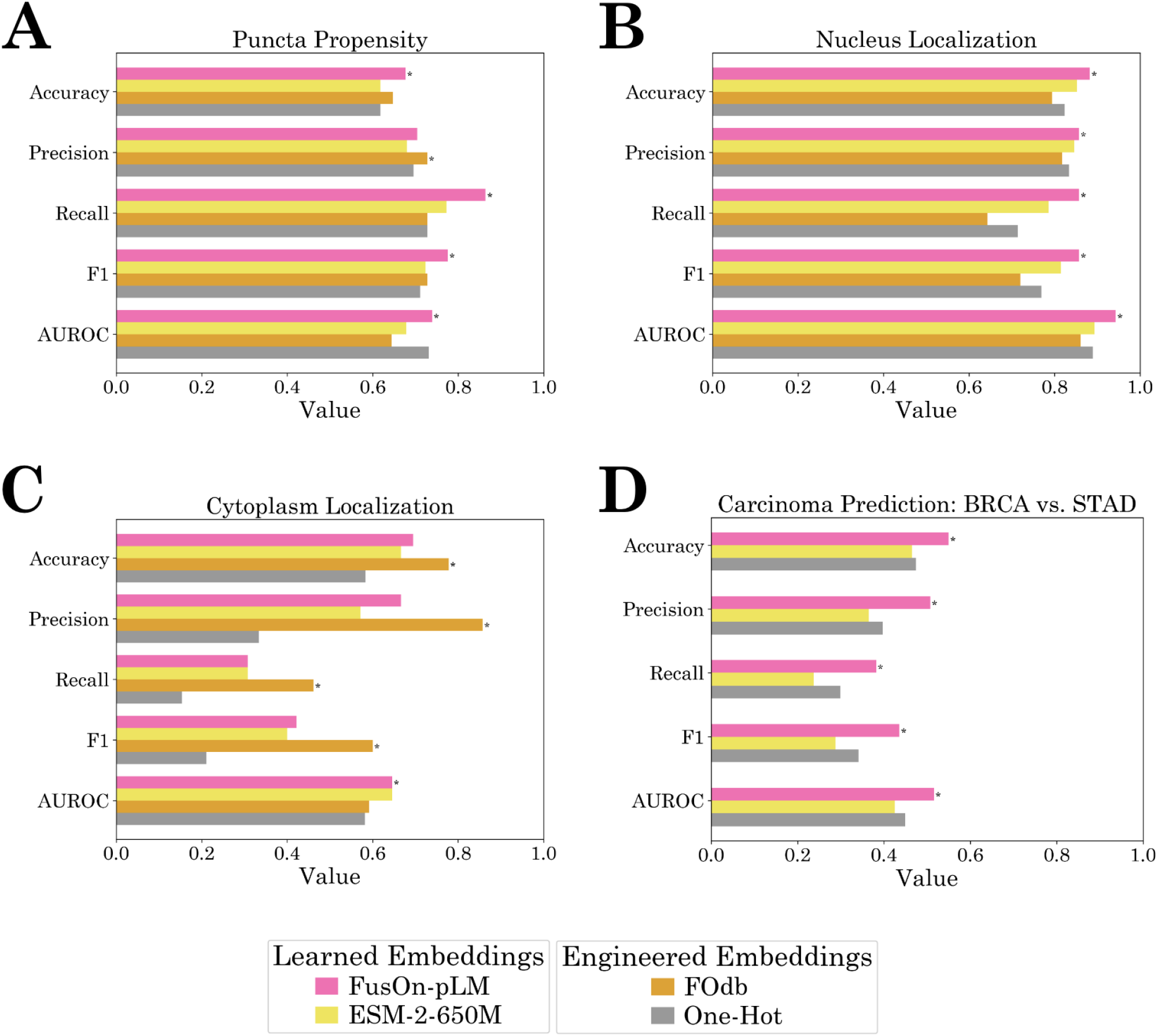
FusOn-pLM embedding benchmarking on fusion oncoprotein-specific tasks. **A-C)** XGBoost binary classifiers utilize FusOn-pLM, ESM-2-650M, FOdb, and one-hot embeddings to predict **A** propensity of puncta formation, **B** puncta localization to the nucleus, and **C** puncta localization to the cytoplasm. **D)** XGBoost binary classifiers utilize FusOn-pLM, ESM-2-650M, and one-hot embeddings to classify fusion oncoproteins as causing BRCA (breast invasive carcinoma) or STAD (stomach adenocarcinoma). FOdb embeddings are not available for these sequences, as they were manually curated for under 200 fusion proteins.

### FusOn-pLM can accurately predict disordered content in wild-type and fusion oncoproteins

Given that fusions are structurally disordered, we hypothesized that FusOn-pLM’s embeddings may encode information pertinent to the properties of intrinsically disordered regions (IDRs), specifically the radius of gyration (*R*_*g*_), defined as the average distance between a protein’s residues and its center of mass, as well as asphericity, which quantifies a protein’s ensemble shape and molecular conformation [Lotthammer et al., 2024]. Comparing FusOn-pLM, ESM-2-650M, and one-hot embeddings against ground truth values of *R*_*g*_ and asphericity, we demonstrate that FusOn-pLM embeddings outperform ESM-2-650M and one-hot embeddings in correctly predicting asphericity (Figure 3A). We also find that FusOn-pLM and ESM-2-650M embeddings achieve equivalent performance and FusOn-pLM strongly outperforms one-hot embeddings when predicting *R*_*g*_ (Figure 3B).

**Figure 3.**
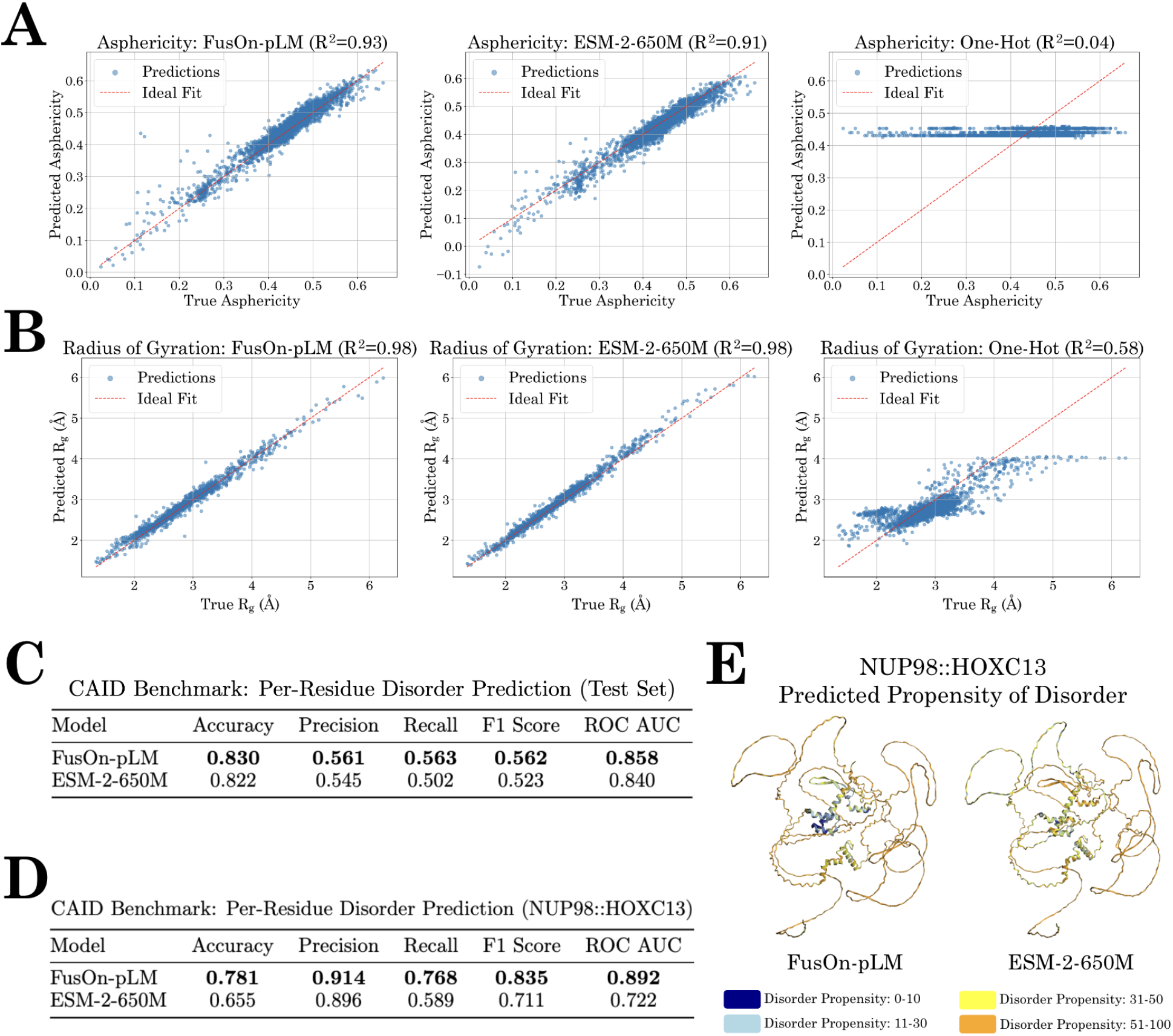
FusOn-pLM prediction of IDR properties and regions. **A-B)** Comparison of FusOn-pLM embeddings with ESM-2-650M embeddings on predicting **A** asphericity and **B** *R*_*g*_ via an MLP head. **C-D)** Classification of disordered residues using FusOn-pLM embeddings and ESM-2 embeddings for **C** CAID test set proteins and **D** fusion oncoprotein NUP98::HOXC13 (which was not included in CAID). **E** Visualization of FusOn-pLM embeddings and ESM-2 embeddings’ predictions of disorder propensity on the AlphaFold3-predicted structure of NUP98::HOXC13. Disorder probabilities are shaded according to the legend for interpolation.

Having proven FusOn-pLM’s ability to predict IDR properties, we sought to compare FusOn-pLM and ESM-2-650M’s ability to identify IDR regions in input protein sequences, via the prediction of per-residue disorder probabilities specified in the CAID dataset [Necci et al., 2021]. Our results demonstrate that FusOn-pLM embeddings achieve stronger performance across all metrics compared to ESM-2-650M (Figure 3C). As such, we then questioned whether FusOn-pLM embeddings could accurately distinguish between structured and disordered residues in fusion oncoproteins. As a demonstration, we focused on NUP98::HOXC13, a known driver of acute myeloid leukemia [Tosić et al., 2009]. We show the FusOn-pLM embedding-based predictions outperform ESM-2-650M embeddings on all classification metrics for this fusion (Figure 3D). Specifically, when visualizing the per-residue disorder probability predictions by differentially coloring disordered and structured residues, we establish that FusOn-pLM correctly identifies the DNA binding domains of NUP98::HOXC13 as structured, coloring them light and dark blue, while ESM-2-650M represents these coils as disordered, coloring them mostly yellow to orange (Figure 3E). Similar patterns are observed for other fusion oncoproteins, including EWSR1::WT1, as well as wild-type proteins, such as SF3B1 (Figure S1). Overall, our results suggest that FusOn-pLM accurately encodes disorder-related information in its embeddings and can thus more effectively represent fusion oncoproteins.

### FusOn-pLM embeddings encode physicochemical features of fusion oncoproteins

The primary objective of FusOn-pLM is to provide feature-rich representations of fusion oncoproteins which are distinct from their head and tail protein counterparts, enabling fusion-specific binder design applications. Given this aim, we visualized FusOn-pLM embeddings in a two-dimensional context to concretely assess the model’s capability in achieving embedding differentiation (Figure 3). Via t-SNE visualization of the generated embeddings, we observe clear and distinct separation between FusOn-pLM fusion embeddings and embeddings of the head and tail proteins for the well-studied fusion oncoproteins PAX3::FOXO1, EWS::FLI1, SS18::SSX1, and EML4::ALK (Figure 4A). The relative distance between the final embeddings suggest that FusOn-pLM learns fusion oncoprotein-specific information in its embeddings that yield distinct, yet accurate, numerical representations of these sequences.

**Figure 4.**
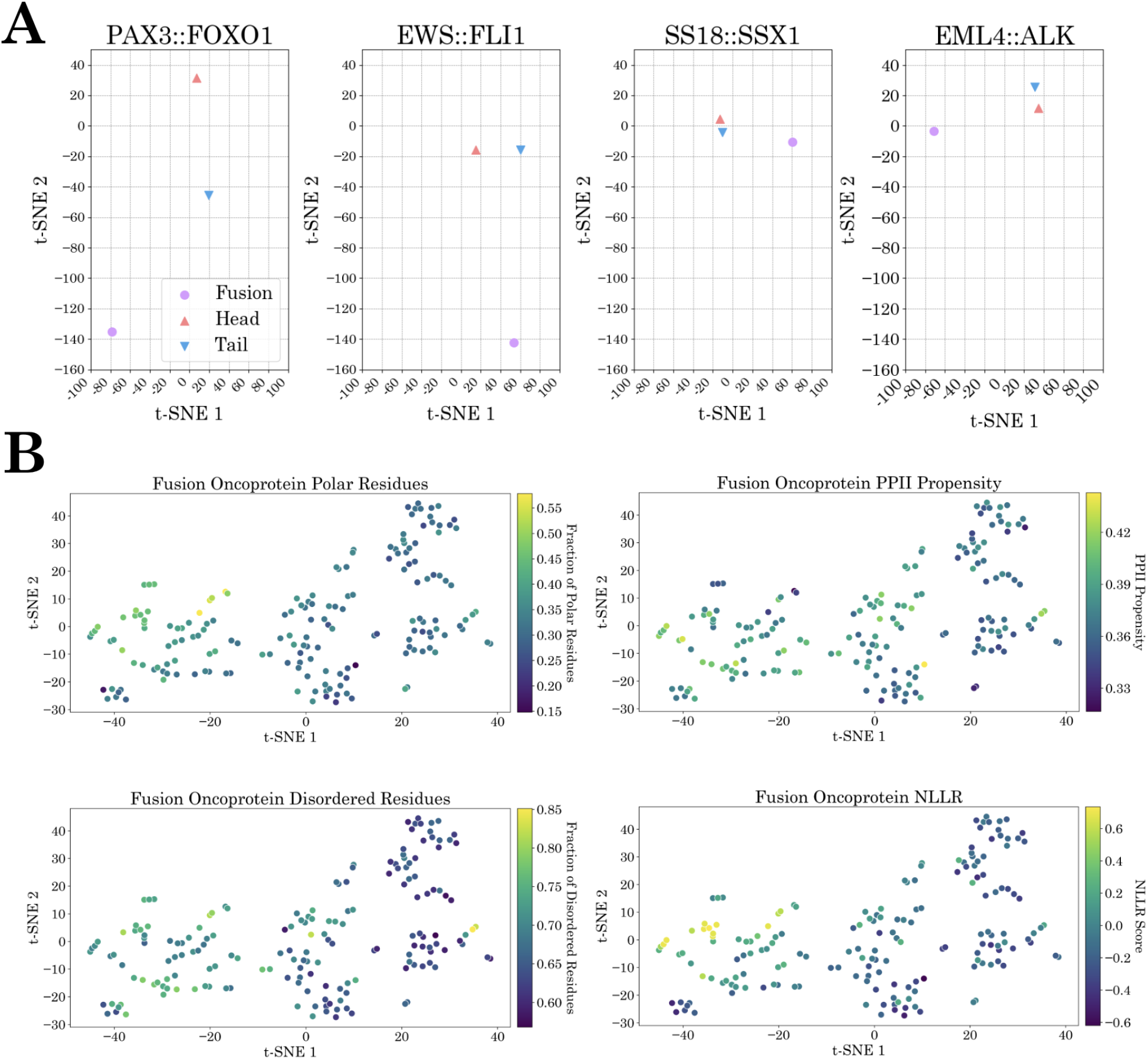
Embedding exploration. **A** FusOn-pLM embedding visualization of fusion oncoproteins, and cognate head and tail protein sequences via t-SNE. Four widely-examined fusion oncoproteins are included (also visualized in Figure 1): PAX3::FOXO1, EWS::FLI1, SS18::SSX1, EML4::ALK. **B** FusOn-pLM embeddings of 177 test set proteins from FOdb, colored by sequence-derived physicochemical properties: fraction of polar residues, fraction of disordered charged residues, propensity to form pi-pi and pi-cation interactions, and NLLR score for prion-like domains.

Finally, we sought to evaluate whether FusOn-pLM embeddings are spatially organized by sequence properties which may drive condensate formation and disease progression. As shown in Figure 4B and Figure S2, FusOn-pLM largely clusters sequences by key properties such as the fraction of polar, charged, and disordered residues as well as the propensity to form pi-pi and pi-cation interactions (quantified as the PScore value)[Vernon et al., 2018] and prion-like domains, via the PLAC NLLR score. [Lancaster et al., 2014] This indicates the semantically expressive and physicochemically-relevant nature of FusOn-pLM embeddings.

## Discussion

In this work, we introduce FusOn-pLM, an ESM-2-based pLM fine-tuned to generate fusion oncoprotein-specific embeddings. To our knowledge, no pLM has explicitly sought to learn unique characteristics of fusion oncoproteins, which differ from most proteins due to their highly disordered nature and altered structural and functional properties driving oncogenic transformation. Our benchmarking results establish that via a binding site-biased MLM training stategy, FusOn-pLM embeddings outperform those of the original ESM-2-650M model [Lin et al., 2023], as well baseline FOdb descriptor embeddings [Tripathi et al., 2023], on fusion oncoprotein-related tasks, and retain distinct representations of fusion proteins from their head and tail counterparts. While FOdb embeddings do perform strongly on certain tasks, such as cytoplasm localization, their inherent static nature precludes their utilization for design tasks via methods such as contrastive learning, autoregressive generation, and diffusion. Additionally, FOdb embeddings were curated through a comprehensive but laborious process for a set of under 200 fusion sequences, hindering their application to the thousands of additional fusion oncoproteins. Finally, we demonstrate that by training on fusion oncoprotein sequences, which represent a large class of IDR-containing proteins, FusOn-pLM embeddings improve upon ESM-2-650M predictions on IDR detection and property prediction tasks.

Recently, our lab has trained ESM-2-based models to generate peptides provided only the sequence of the target protein, facilitating the design of CRISPR-like peptide-E3 ubiquitin ligase fusions for the target-specific proteasomal degradation of conformationally-diverse protein substrates [Bhat et al., 2023, Chen et al., 2023]. As such, our next steps will be to replace ESM-2 embeddings in these models with FusOn-pLM embeddings, enabling fusion-specific degrader design. Since post-translational modifications (PTMs) are also well known to affect the oncogenic activity of fusion oncoproteins [Yu et al., 2023, Thalhammer et al., 2015, Pan and Chen, 2022], we plan to retrain FusOn-pLM with our recent PTM-Mamba pLM [Peng et al., 2024], which effectively tokenizes PTMs, enabling both fusion- and PTM-specific therapeutic design. Finally, by leveraging recent advancements in gene delivery, such as lipid nanoparticles (LNPs) and adeno-associated viral (AAV) vectors [Hou et al., 2021, Wang et al., 2024], we envision that fusion-specific biologics may eventually serve as safe and efficacious therapeutics for fusion-positive cancer patients. Overall, the results of our study motivate the use of FusOn-pLM embeddings for downstream fusion oncoprotein design tasks, serving as a major step toward this goal.

## Declarations

## Acknowledgements

We thank Mark III Systems and the Duke Computing Cluster for computing support. We further thank Zhangzhi Peng, Yinuo Zhang, and Tianlai Chen for their insights related to the manuscript. The work was supported by the National Cancer Institute (Award #R21CA278468), the Wallace H. Coulter Foundation, and The Hartwell Foundation.

## Author Contributions

S.V. designed and implemented masking strategies and trained FusOn-pLM. S.V., S.G., K.K., R.P., and P.V. performed model benchmarking and visualizations. S.V., S.G., K.K., and P.C. wrote and reviewed the manuscript. P.C. conceived, designed, directed, and supervised the study.

## Data and Materials Availability

All data needed to evaluate the conclusions are presented in the paper and tables. Model weights and training code will be released at https://huggingface.co/ChatterjeeLab/FusOn-pLM.

## Supplementary Information

**Figure S1:**
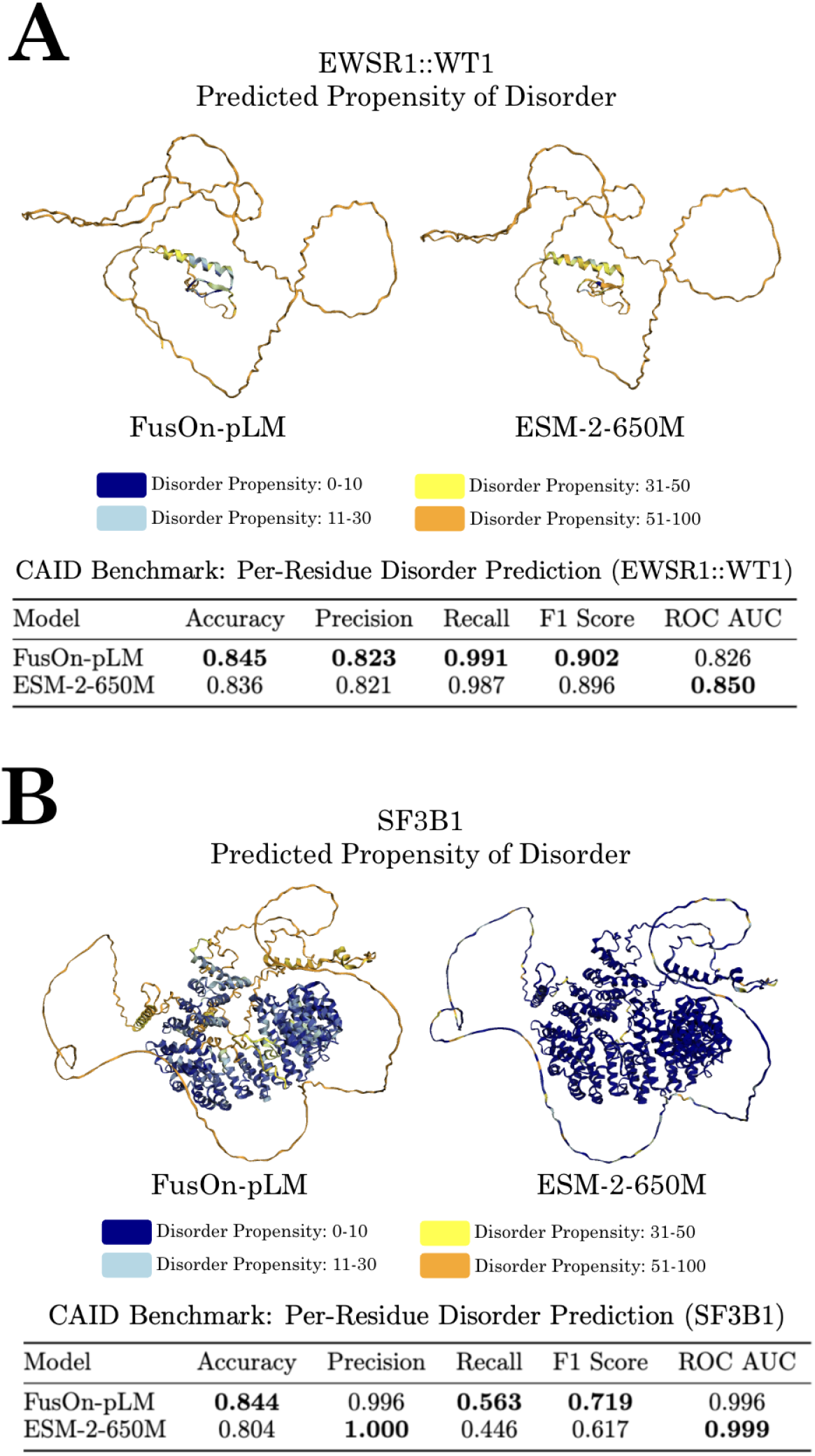
IDR prediction and visualization. Disordered residues were classified using FusOn-pLM and ESM-2-650M embeddings and were vizualized as predicted disordered propensities on the AlphaFold3-predicted structure of **A** fusion oncoprotein EWSR1::WT1 (which was not included in the CAID test set) and **B** CAID test set protein SF3B1. Disorder propensities are shaded according to the legend for interpolation.

**Figure S2:**
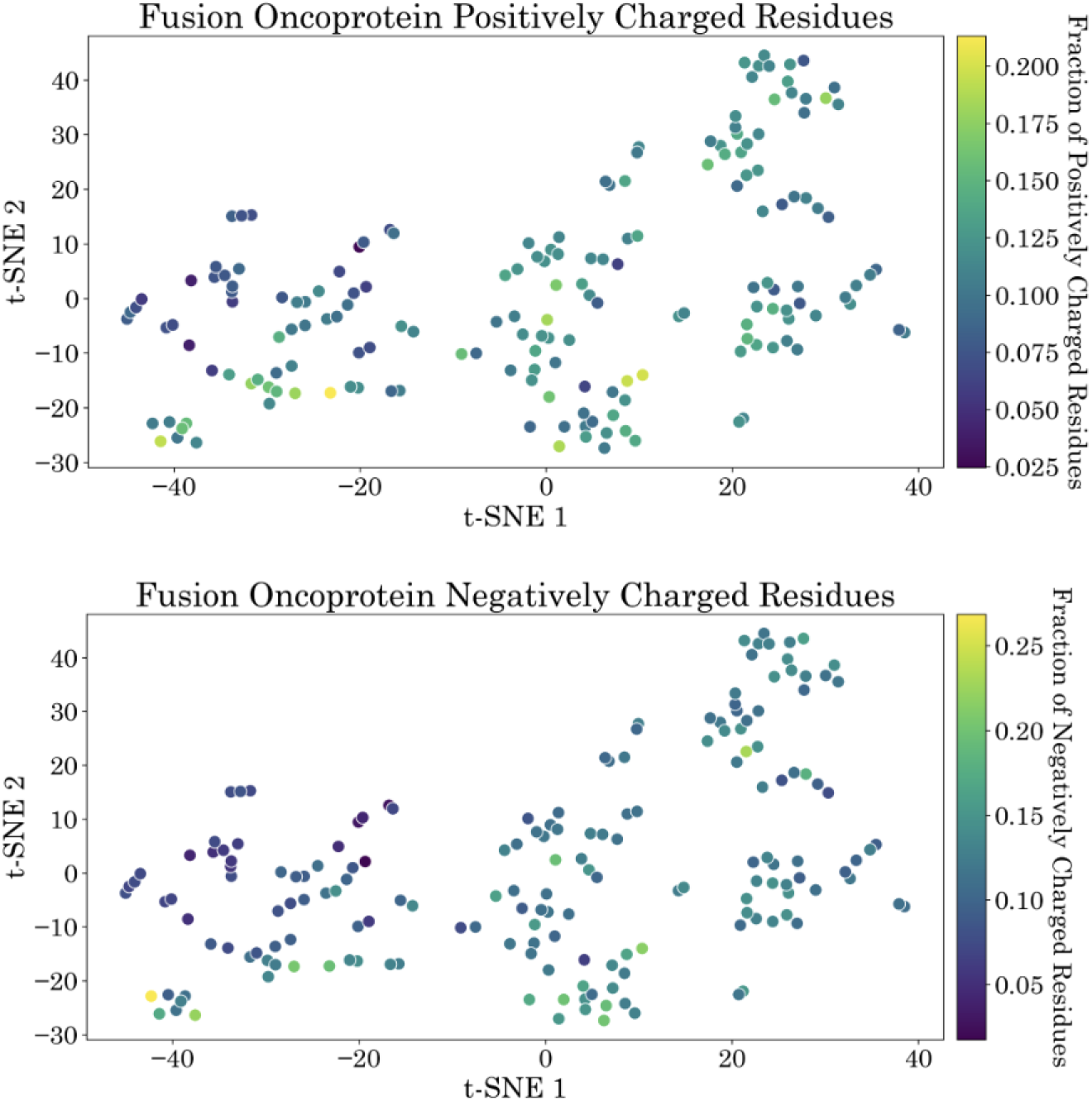
Embedding exploration. FusOn-pLM embeddings of 177 test set proteins from FOdb, colored by sequence-derived physicochemical properties: fraction of positively charged residues and fraction of negatively charged residues.

